# Infant RSV infection desensitizes β2-adrenergic receptor via CXCL11-CXCR7 signaling in airway smooth muscle

**DOI:** 10.1101/2025.01.13.632772

**Authors:** Caiqi Zhao, Alice E. Taliento, Elise M Belkin, Rachel Fearns, Paul H. Lerou, Xingbin Ai, Yan Bai

## Abstract

**Rationale:** Airflow obstruction refractory to β2 adrenergic receptor (β2AR) agonists is an important clinical feature of infant respiratory syncytial virus (RSV) bronchiolitis, with limited treatment options. This resistance is often linked to poor drug delivery and potential viral infection of airway smooth muscle cells (ASMCs). Whether RSV inflammation causes β2AR desensitization in infant ASMCs is unknown.

**Objectives:** To investigate the interaction of RSV inflammation with the β2AR signaling pathway in infant ASMCs

**Methods:** Infant precision-cut lung slices (PCLSs) and mouse pup models of RSV infection were subjected to airway physiological assays. Virus-free, conditioned media from RSV-infected infant bronchial epithelial cells in air-liquid interface (ALI) culture and nasopharyngeal aspirates (NPA) from infants with severe RSV bronchiolitis were collected and applied to infant PCLSs and ASMCs. Cytokines in these samples were profiled and assessed for the effects on β2AR expression, cell surface distribution, and relaxant function in ASMCs.

**Measurements and Main Results:** Conditioned media and NPA induced similar resistance to β2AR agonists in ASMCs as RSV infection. Cytokine profiling identified CXCL11 as one of the most elevated signals following RSV infection. CXCL11 activated its receptor CXCR7 in a complex with β2AR in ASMCs to promote β2AR phosphorylation, internalization, and degradation. Blockade of CXCR7 partially restored airway relaxation in response to β2AR agonists in infant PCLSs and mouse pup models of RSV infection.

**Conclusions:** The CXCL11-CXCR7 pathway plays a critical role in β2AR desensitization in ASMCs during RSV infection and represents a potential therapeutic target in alleviating airflow obstruction in infant RSV bronchiolitis.

## INTRODUCTION

RSV infection is a leading cause of acute lower respiratory tract inflammation in infants and young children (1, 2). To date, the development of RSV vaccine in infants has been unsuccessful. Current strategies to protect infants from RSV infection include vaccination during late pregnancy and postnatal administration of two FDA-approved antibodies, Palivizumab and Nirsevimab, that are effective in reducing RSV-related hospitalization (3, 4). However, these prophylactic approaches do not offer 100% protection. For infants with severe RSV infection, the mainstay of treatment remains limited to supportive care (5–7). A devastating clinical symptom is airflow obstruction refractory to β2AR agonists. Most recent evidence-based clinical practice guidelines do not recommend bronchodilators (6, 7), leaving no drug available to alleviate airflow obstruction in infant RSV bronchiolitis. A prevailing view in the field attributes intractable airflow obstruction in infant RSV disease to profound epithelial sloughing and mucus hyperplasia that cause airway plugging and consequentially limit the accessibility of inhaled β2AR agonists to the affected area (8, 9). RSV was also shown to infect ASMCs *in vitro* to reduce β2AR levels (10, 11). However, lung histology from fatal cases of infant RSV disease demonstrates little evidence for direct RSV infection of ASMCs *in vivo*(12–14). Therefore, while limited accessibility of β2AR agonists remains a possibility, additional factors, such as inflammation in infant RSV bronchiolitis, may also play a role in impaired β2AR agonist responses in ASMCs. Due to technical difficulties in accessing diseased airways in infants with RSV bronchiolitis, whether and how inflammation dysregulates ASMC contraction and relaxation in infant RSV disease is completely unknown (15).

β2AR agonists relax the airway by activating the cyclic AMP (cAMP)-protein kinase A (PKA) pathway to suppress Ca^2+^ signaling and reduce Ca^2+^ sensitivity of the contractile apparatus in ASMCs(16, 17). β2AR desensitization is a major mechanism that dampens the relaxant effect of β2AR agonists. This process is initiated by β2AR phosphorylation typically by G protein-coupled receptor (GPCR) kinases (GRKs) or protein kinase A or C (PKA or PKC) (18–20) followed by β-arrestin2-mediated receptor internalization. In respiratory diseases, such as asthma, multiple inflammatory cytokines are known to cause β2AR desensitization in ASMCs, including IL-1β and TNFα that act by triggering PGE2-EP2 dependent PKA activation (21–23). A cAMP-independent mechanism may also contribute to this process. For instance, IL-13 was shown to reduce airway relaxation in response to β2AR agonists by activating ERK (24–26) and interacting with K^+^ channels (27). The established roles of cytokines in β2AR desensitization in asthma suggest that similar communication between inflammation and ASMCs may occur in infant RSV bronchiolitis. Indeed, a previous study showed that RSV infection in BALB/c mice elevated the level of keratinocyte cytokine (the murine homolog of human CXCL8/IL-8) to inhibit the β2AR-mediated ASMC relaxation response (28).

To investigate the role of RSV-induced inflammation in β2AR desensitization in ASMCs, we employed three infant RSV models that have been recently established in our lab (29). These include RSV infection of infant bronchial epithelial cells in ALI as an *in vitro* model, PCLSs prepared from infant donor lungs as an *ex vivo* model, and neonatal mice infected with a modified RSV-A19 stain as an *in vivo* model. These models recapitulate salient clinical features of severe RSV bronchiolitis in infants, including epithelial cell death and airway contraction refractory to β2AR agonists. To translate our findings from these three models into clinical significance, nasopharyngeal aspirates (NPA) from infants who were critically ill from RSV infection were collected as a surrogate to inflammation *in vivo*. Our study assessed changes in cell surface localization and the activity of β2AR in ASMCs following RSV infection and profiled inflammation by cytokine array. In addition, leveraging the *Cxcl11* genetic background difference between Balb/c (*Cxcl11^+/+^)* and C57BL/6 (*Cxcl1^-/-^)* mouse strains, we performed complementary, functional blockade and rescue assays *in vivo*. Our findings have identified CXCL11-CXCR7 as a critical mediator of the communication between airway inflammation and β2AR desensitization in ASMCs in infant RSV bronchiolitis.

## Materials and methods

### Human donor lungs

Lungs from de-identified human donors who were declined for transplantation and had no known airway diseases were purchased from IIAM (International Institute for the Advancement of Medicine). Four infant lungs (2 females and 2 males, aged between 6 days and 16 months) were used in the study to prepare PCLSs and isolate primary ASMCs. This work is deemed nonhuman subject research by the Institutional Review Board (IRB) at Massachusetts General Hospital (MGH).

### Human NPA samples

NPA samples from pediatric patients with bronchiolitis were collected under an approved IRB protocol (No. 2019P003296; PI: Lerou). For this study, NPA samples from 6 RSV-infected patients (2 weeks to 24 weeks), 2 rhinovirus-infected patients (11 weeks and 36 weeks), and one patient infected with metapneumovirus (28 weeks) were used.

### Mouse model of RSV infection

BALB/c and C57BL/6 mice were purchased from the Jackson Laboratory, USA. All procedures in mice were approved by the Institutional Animal Care and Use Committee (2019N000137, PI: X.A). Mouse pups were intranasally inoculated with medium or RSV A2-19 (1×10^6^ PFU (plaque-forming unit), diluted in 10 µL medium) on postnatal days 7 and 8. Treatment with CXCL11 (200 ng/ml, R&D, 672-IT-025) or CXCR7 antagonist (10 μM, MCE, HY-139643) was performed by mixing RSV and the reagent in a volume of 10 µL following by intranasal inoculation. Mice were euthanized on postnatal day 10 and lungs were harvested for multiple assays.

### Statistical Analysis

Details of the number of samples and experiments were presented in the corresponding figure legends and result sections. Data were analyzed using GraphPad Prism 10 (San Diego, CA) and reported as mean ± SEM. For statistical analysis of non-parametric data in most experiments, the Wilcoxon test was applied to compare paired measurements of the same sample under two experimental conditions, the Mann– Whitney U test was applied to compare non-paired, two experimental conditions, the Kruskal-Wallis test with the post-hoc analysis using Benjamini, Krieger, or Yekutieli (BKY) methods was applied to compare multiple conditions. For statistical analysis of parametric data with one or two independent variables, a one-way or two-way ANOVA test with the post-hoc Tukey or Sidak test was used. p < 0.05 was considered statistically significant. Detailed description of the materials and the methods used in the study, including RSV preparation and infection, human PCLS preparation, primary human ASMC culture, epithelial ALI culture, airway contraction and relaxation assays, cytokine and protein analyses, and gene expression assays are available in online supplementary documents.

## RESULTS

### RSV-infected infant PCLSs show dampened airway relaxation responses to β2AR agonists

To assess whether infant RSV infection affects airway smooth muscle relaxation in response to β2AR agonists, we employed the infant PCLS model of RSV infection. RSV (A2 strain, 1×10^6^ PFU) was incubated with PCLSs prepared from de-identified infant (0-18 months) donor lungs for 3 days to establish infection (Fig. 1A). Double staining for RSV F protein and smooth muscle α actin (αSMA) detected the virus almost exclusively in the bronchial epithelial layer, consistent with highly selective ciliated epithelial cell tropism of RSV (13, 30, 31) (Fig. 1B). A few RSV F^+^ cells scattered in the parenchyma were likely phagocytes (Fig. 1B). No RSV was found in αSMA^+^ ASMCs, as previously shown by pathology of infant lungs with fatal RSV infection (12, 13). Prior to the contraction-relaxation assay, epithelial debris and mucus in the airway lumen were cleared by gently flushing PCLSs with the culture medium to prevent interference with airway responses. Compared to mock control, RSV-infected infant PCLSs exhibited enhanced airway contraction in response to histamine at doses higher than 0.2 µM (Fig. 1C, 1D). Once reaching maximum contraction at 10 µM histamine, airway relaxation in response to formoterol, a long-acting β2AR agonist, was profoundly blunted in RSV-infected infant PCLSs compared to ∼65% relaxation in mock control (Fig. 1D, 1E). We then tested whether the blunted relaxation response was secondary to exaggerated airway contraction. At 0.2 μM histamine, a concentration that induced comparable ∼30% airway contraction in both groups (Fig. 1D), formoterol remained less effective in RSV-infected infant PCLSs compared to mock control (Fig. 1F). Therefore, the infant PCLS model reproduces impaired airway relaxation responses to β2AR agonists found in infant RSV bronchiolitis.

**Fig 1.**
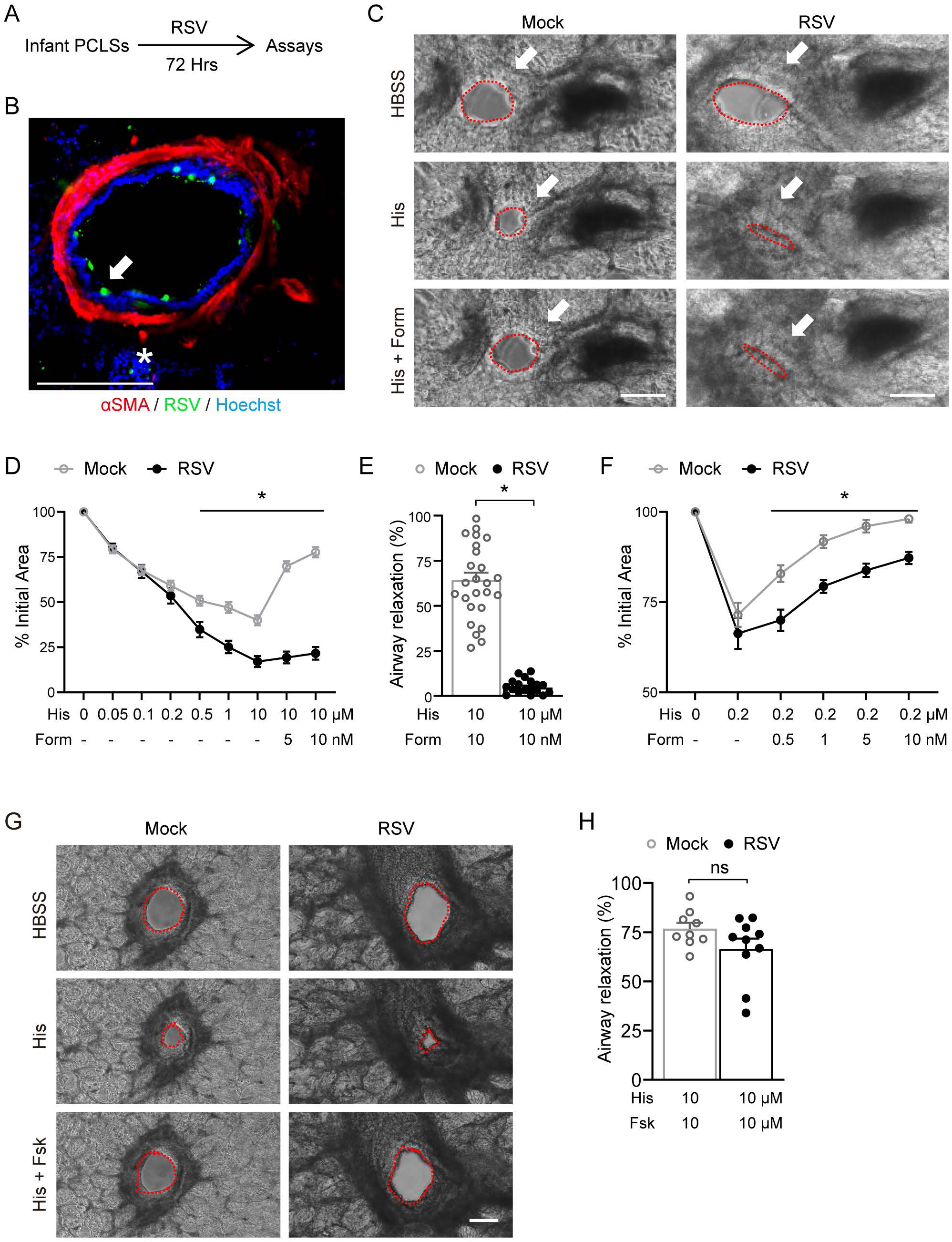
RSV infection of infant PCLSs enhanced airway contraction and blunted relaxation responses to β2AR agonists. (**A**) Schematic of RSV A2 (1×10^6^ PFU) infection of PCLSs prepared from infant lungs (n=4 donors). After 72 hours, infant PCLSs were subjected to contraction and relaxation assays. (**B**) A representative confocal image of double fluorescence staining for RSV F protein and αSMA in RSV-infected infant PCLSs. Arrow marks RSV F^+^ epithelial cells. Asterisk marks rare non-epithelial RSV F^+^ cells. (**C**) Representative bright-field images of airways in Mock- and RSV-infected infant PCLSs at baseline, in response to 10 μM histamine (His), and then His +10 nM Formoterol (Form). (**D**) Summary plots showing airway contraction in response to His and relaxation in response to Form at indicated concentrations in Mock- and RSV-infected infant PCLSs. The contraction was presented as % of the airway lumen size at baseline. N=20-22 PCLSs from 4 donors. (**E**) Bar graph showing relaxation responses of 10 µM His-precontracted airways to 10 nM Form in Mock- and RSV-infected infant PCLSs. Relaxation was calculated as % of His-reduced luminal area that was recovered by Form. Each mark represents one airway. (**F**) Summary plots showing airway contraction by 0.2 μM His and relaxation in response to Form at indicated concentrations in Mock- or RSV-infected infant PCLSs. N=8-9 PCLSs from 4 donors. (**G**) Representative bright field images of airways in Mock- and RSV-infected infant PCLSs at baseline and in response to 10 μM His and then His+10 μM Forskolin (Fsk). (**H**) Bar graph showing relaxation of 10 µM His-precontracted airways in response to 10 μM Fsk. Relaxation was calculated as % of His-reduced luminal area that was recovered by Fsk. Each mark represents one airway. Data represent mean ± SEM. The airway lumen was outlined by a dotted line. Statistical significance was calculated by the two-way ANOVA test with a post-hoc Sidak test in (D) and (F) and the Mann-Whitney test in (**E**) and (H). *p<0.05. ns, not significant. Scale bars, 100 μm.

Since β2AR agonists act through the cAMP-mediated pathway, we tested whether downstream cAMP signaling and the relaxation machinery in infant ASMCs were disrupted by epithelial RSV infection. To do so, we treated infant PCLSs with forskolin (FSK), a cell-permeable compound that can directly activate the adenylate cyclase-cAMP pathway to induce airway relaxation. FSK treatment elicited similar levels of airway relaxation in mock and RSV-infected infant PCLSs (Fig. 1G, 1H). These findings indicate that infant RSV infection of ciliated cells can suppress ASMC relaxation by disrupting components specific to the β2AR pathway.

### Epithelial inflammation following infant RSV infection induces β2AR desensitization in ASMCs

Since the infant PCLS model has no recruited immune cells, inflammatory signals from RSV-infected bronchial epithelium may play a major role in the dysregulation of airway contraction and relaxation. To test this hypothesis, we employed the *in vitro* model of RSV infection (4×10^5^ PFU) in ALI cultures of neonatal bronchial epithelial cells that were differentiated from tracheal aspirate-derived BSCs as previously described (29, 32). At day 2 post-infection, conditioned medium (CM) in the bottom chamber of ALI culture was collected and used to treat infant PCLSs (Fig. 2A). Because RSV virions are released only from the apical surface of the infected ciliated cells, no RSV was found in CM (29). After treatment for 48 hours, compared to Mock-CM, RSV-CM induced airway hypercontraction in response to histamine and blunted relaxation responses to formoterol in infant PCLSs (Fig. 2B), similar to the findings in the infant PCLS model of RSV infection (Fig. 1C-F). Therefore, secreted signals from RSV-infected infant bronchial epithelial cells are sufficient to dysregulate airway contraction and relaxation.

**Fig 2.**
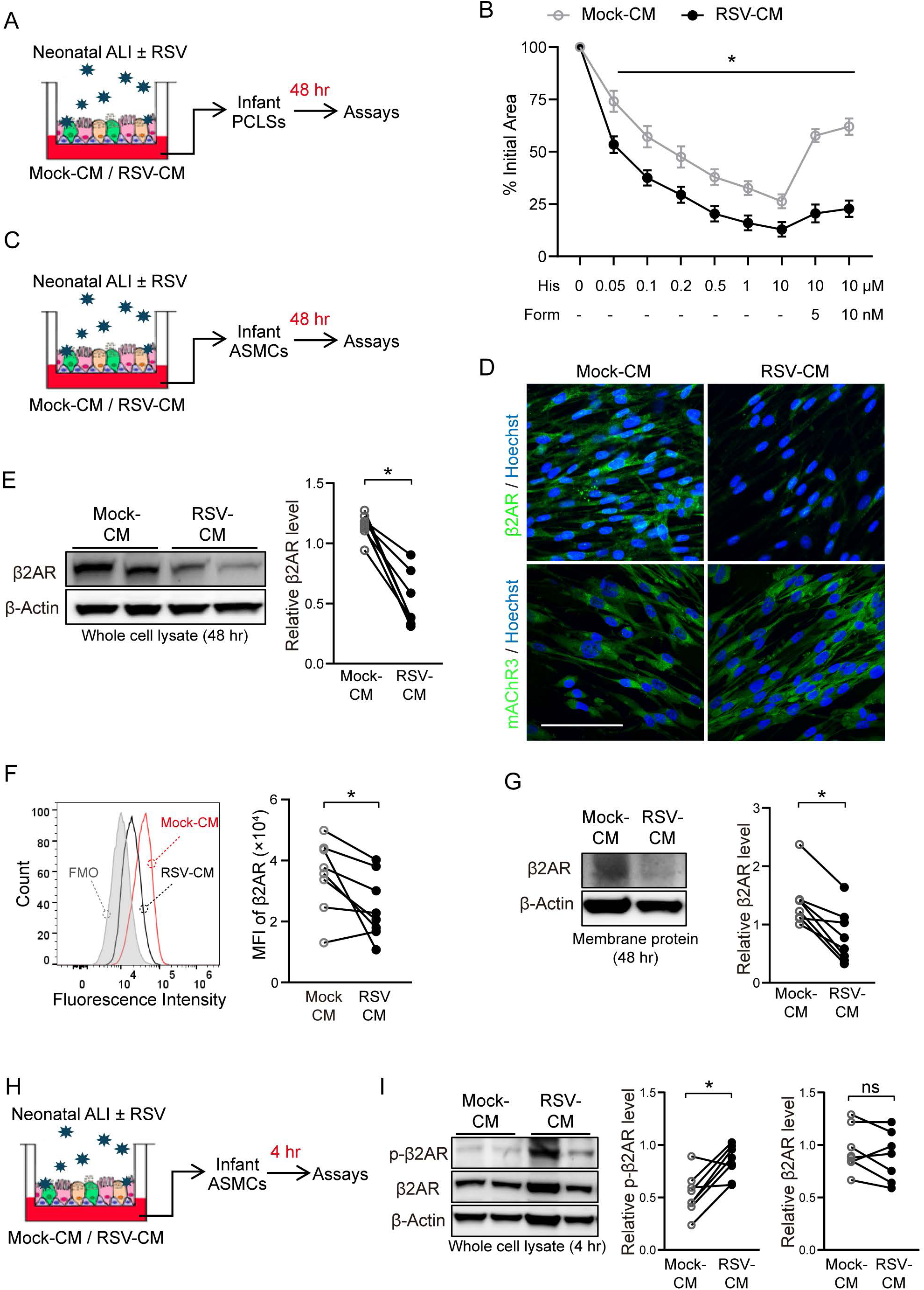
RSV-induced epithelial inflammation is sufficient to cause β2AR desensitization in ASMCs. (**A**) Schematic of conditioned medium (CM) collection from Mock- and RSV-infected (MOI 2, 4×10^5^PFU) neonatal epithelial cells in ALI on day 2 post-infection followed by treatment of infant PCLSs with CM for 48 hours. (**B**) Summary plots of airway contraction in response to His and relaxation in response to Form at indicated concentrations in Mock- and RSV-CM treated infant PCLSs. The contraction was measured as % of the airway lumen size at baseline. N=9-10 infant PCLSs from 4 donors. (**C**) Schematic of Mock- and RSV-CM treatment of primary infant ASMCs for 48 hours followed by assays for cell surface and total levels of β2AR in (D-G). (**D**) Representative fluorescence images showing antibody staining for β2AR and muscarinic acetylcholine 3 receptor (mAChR3) in primary infant ASMCs following treatment with Mock- or RSV-CM. Nuclei were labelled by Hoechst dye. Scale bars, 100 μm. (**E**) Representative Western blot for β2AR in Mock- and RSV-CM-treated infant ASMCs. β-actin was the loading control in densitometry. N=4 donors with 2 independent repeats. (**F**) Representative flow cytometry plots showing median fluorescence intensity (MFI) of cell surface β2AR in infant ASMCs. Fluorescence minus one (FMO) control was included. N=4 donors with 2 independent repeats. (**G**) Representative Western blot for biotinylated cell surface β2AR. Biotin crosslinking of membrane proteins was performed using infant ASMCs. Biotinylated proteins were enriched by neutravidin agarose resins prior to Western blot. β-actin in whole cell lysate was loading control in densitometry. N=4 donors with 2 independent repeats. (**H**) Schematic of Mock- and RSV-CM treatment of primary infant ASMCs for 4 hours followed by Western blot assay for phosphorylated β2AR (p-β2AR) in (I). (**I**) Representative Western blot of p-β2AR. β-actin was the loading control in densitometry. N=4 donors with 2 independent repeats. Data in (B) represent mean ± SEM. Statistical significance was calculated by a two-way ANOVA test with a post-hoc Sidak test in (B) and by a Wilcoxon test in (E)-(I). *p<0.05.

We chose to focus on how RSV infection blunts the airway relaxation response to β2AR agonists, as intractable airflow obstruction is a leading cause of morbidity and mortality in infant RSV bronchiolitis. We first profiled infant ASMCs from 3 different donors for global changes in gene expression following RSV-CM treatment by bulk RNA-seq. We identified a total of 146 differentially expressed genes (p<0.05) that were enriched in metabolic and inflammatory pathways (Fig. S1B and D). However, genes involved in the β2AR signaling pathway had no significant change in expression (Fig. S1C). Therefore, the β2AR pathway components may be regulated at post-transcriptional levels by epithelial RSV infection. Since β2AR desensitization via receptor internalization and degradation is a well-established post-translational mechanism, we then assessed changes in the protein level and cell surface localization of β2AR in infant ASMCs following CM treatment for 48 hours (Fig. 2C). Antibody staining for β2AR in fixed ASMCs and Western blot using whole cell lysates detected a significant decrease in the level of β2AR following 48-hour RSV-CM treatment (Fig. 2D, 2E). In contrast, the level of muscarinic acetylcholine receptor 3 (mAChR3) was not affected by RSV-CM (Fig. 2D). To selectively detect cell surface β2AR, we performed two different assays. The first assay was to label β2AR with a specific antibody using live ASMC cultures so that the cell membrane is intact and prevents the antibody from entering cells. After live staining for β2AR, flow cytometry detected lower surface fluorescence in RSV-CM-treated ASMCs than in Mock-CM control (Fig. 2F). The second assay employed biotin crosslinking of membrane proteins followed by avidin-beads enrichment and yielded similar findings of reduced cell surface β2AR following RSV-CM treatment (Fig. 2G). Furthermore, we assessed β2AR phosphorylation (p-β2AR) in ASMCs at 4 hours post-treatment with RSV-CM (Fig. 2H). Western blot showed that RSV-CM-treated infant ASMCs had a significant increase in the level of p-β2AR, while total β2AR levels remained unchanged at this early time point (Fig. 2I). Together, RSV-induced epithelial inflammation in infant lungs is sufficient to cause β2AR phosphorylation, internalization, and degradation in ASMCs, which likely renders desensitized airway relaxation in response to β2AR agonists.

### RSV infection triggered epithelial CXCL11 release to lead β2AR deregulation in infant ASMCs

To identify RSV-induced epithelial inflammatory factors that caused β2AR desensitization in infant ASMCs, we profiled cytokines in CM collected from Mock and RSV-infected neonatal ALI cultures. To narrow down neonatal-specific candidate cytokines, we included cytokine profiles from RSV-infected adult ALIs for comparison (Fig. S2A and B, Fig. 3A). We showed previously that adult bronchial epithelial cells are resistant to severe RSV infection(29). In addition, the RSV-CM from adult ALIs had no effect on infant airway contraction and relaxation (Fig. S2C, S2D). We found that RSV infection caused a greater increase in CXCL11, CXCL10, CXCL9, IL-18Bpa, G-CSF, and IL-19 levels in the CM from neonatal than adult ALI cultures, among which CXCL11 was top-ranked and elevated only in RSV-infected neonatal ALIs (Mock-CM 1.39 ± 0.23 vs RSV-CM 127.53 ± 24.69 (ng per g of total protein), p<0.05), but not in adult ALIs (Mock-CM 0.69 ± 0.06 vs RSV-CM 1.34 ± 0.23 (ng per g of total protein) (Fig. 3A, 3B). These findings identified CXCL11 as a candidate cytokine in triggering β2AR-desensitization in ASMCs following RSV infection of infant bronchial epithelium.

**Fig 3.**
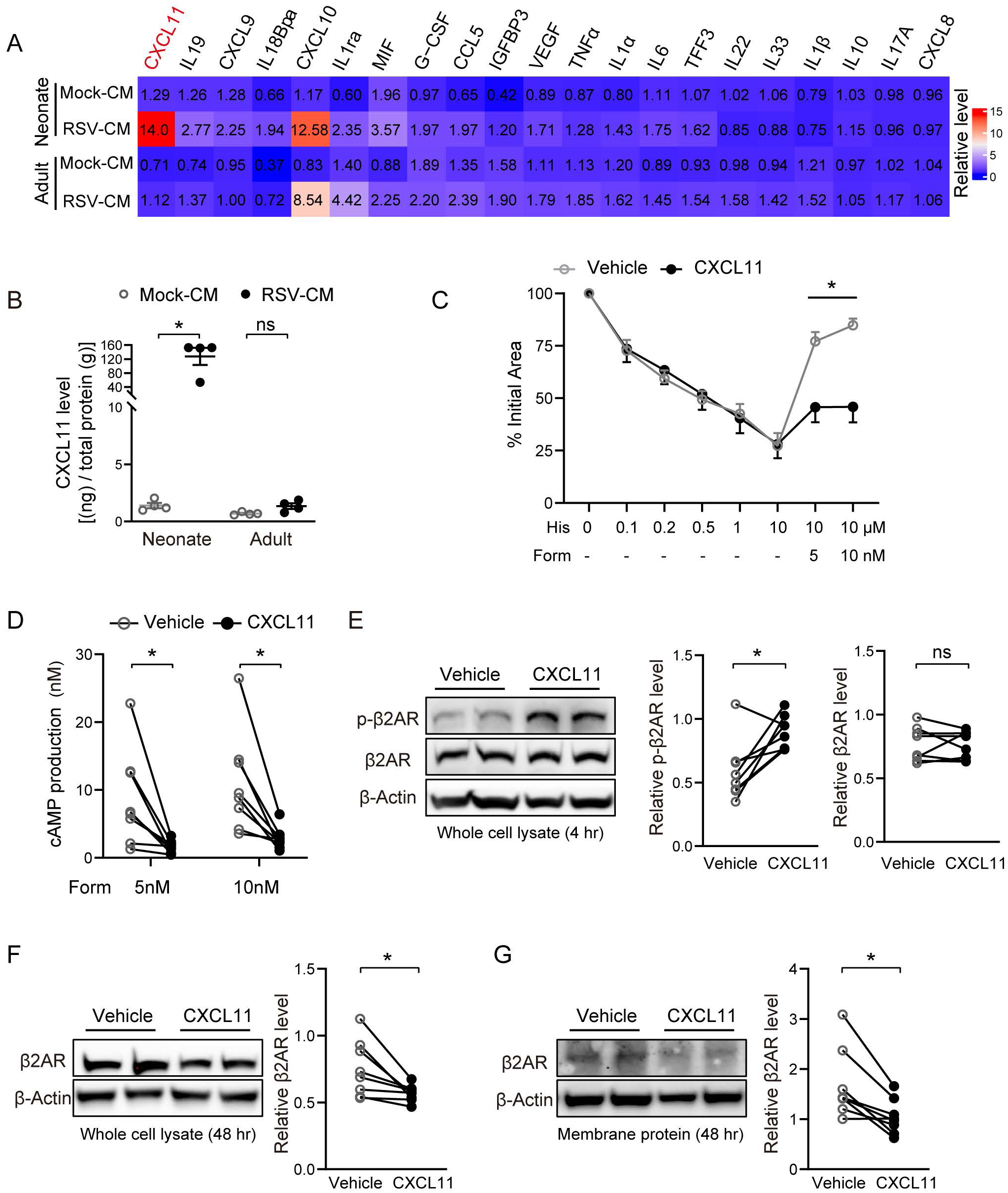
CXCL11 is most elevated in RSV-CM from neonatal ALIs and causes β2AR desensitization in infant ASMCs. (**A**) Heatmap of elevated cytokines in CM collected from Mock or RSV-infected, neonatal and adult ALI cultures by cytokine array. (**B**) ELISA of CXCL11 in Mock- and RSV-CM from neonatal and adult ALI cultures. Data shown were normalized to the total protein level in CM. Each mark represents one donor. (**C**) Summary plots of airway contraction in response to His and relaxation in response to Form at indicated concentrations in infant PCLSs after 48-hr treatment with vehicle or 200 ng/ml CXCL11. N=8 PCLSs from 4 donors for the vehicle group. N=9 infant PCLSs from 4 donors in the CXCL11 group. (**D**) Form-induced cAMP production in infant ASMCs that were pre-treated with vehicle or CXCL11 (200 ng/ml) for 48 hours. N=4 donors with 2 independent repeats. (**E**) Representative Western blot for p-β2AR and total β2AR infant ASMCs after 4-hr treatment with vehicle or CXCL11 (200 ng/ml). β-actin is the loading control in densitometry. N=4 donors with 2 independent repeats. (**F** and **G**) Representative Western blot for total (**F**) and cell surface β2AR (**G)** in infant ASMCs after 48-hr treatment with solvent or 200 ng/ml CXCL11. Cell surface β2AR was crosslinked biotin and then enriched by neutravidin agarose resins. N=4 donors with 2 independent repeats. Statistical significance was calculated by the Mann-Whitney test in (B), a two-way ANOVA test with a post-hoc Sidak test in (C), and a Wilcoxon test in (D)-(G) *p<0.05. ns, not significant.

To test the role of CXCL11 in desensitized ASMC responses to β2AR agonists, we treated infant PCLSs with CXCL11 (200 ng/ml) for 48 hours, followed by airway contraction and relaxation assays (Fig. 3C). CXCL11 had no effect on airway contraction in response to Histamine. However, CXCL11 treatment blunted airway relaxation in response to formoterol (Fig. 3C) as RSV (Fig. 1D) and RSV-CM (Fig. 2B) did. Consistent with the β2AR desensitization effect, 48-hour CXCL11 treatment significantly reduced formoterol-induced cAMP production in ASMCs (Fig. 3D), the total level of β2AR in cell lysates (Fig. 3F), and the cell surface level of β2AR (Fig. 3G). In addition, 4-hour CXCL11 treatment induced β2AR phosphorylation but had no effect on the total receptor level (Fig. 3E). Together, RSV infection of infant epithelial cells activates the expression and secretion of CXCL11 and CXCL11 can signal to ASMCs to downregulate β2AR and blunt relaxation responses to β2AR agonists.

### CXCL11 interacts with CXCR7 on infant ASMCs to cause β2AR desensitization

CXCL11 has two known receptors, CXCR3 and CXCR7. Bulk RNA-seq of human ASMCs (Fig. S1) showed that *CXCR3* mRNA levels were almost undetectable. In comparison, *CXCR7* was robustly expressed with 2^10^ fold more counts. Consistent with the results of mRNA analyses, CXCR7 was readily detectable in ASMCs by antibody staining and Western blot (Fig. 4A and 4B). In addition, in contrary to β2AR that was downregulated following RSV-CM treatment (Fig. 2C-G), the level of CXCR7 in infant ASMCs was maintained regardless of CM treatment (Fig. 4A and B). To assess the role of CXCR7 as the receptor of CXCL11 in mediating β2AR desensitization following RSV-CM treatment of infant PCLSs, we used a neutralizing antibody for CXCR7 in the RSV-CM treatment experiment. We found that CXCR7 neutralization rescued the relaxant effect of formoterol to a level similar to the response in Mock-CM-treated infant PCLSs (Fig. 4C). In addition, CXCR7 neutralization prevented β2AR downregulation in infant ASMCs following treatment with RSV-CM (Fig. 4D). These findings indicate that activation of the CXCL11-CXCR7 axis in ASMCs promotes β2AR desensitization.

**Fig 4.**
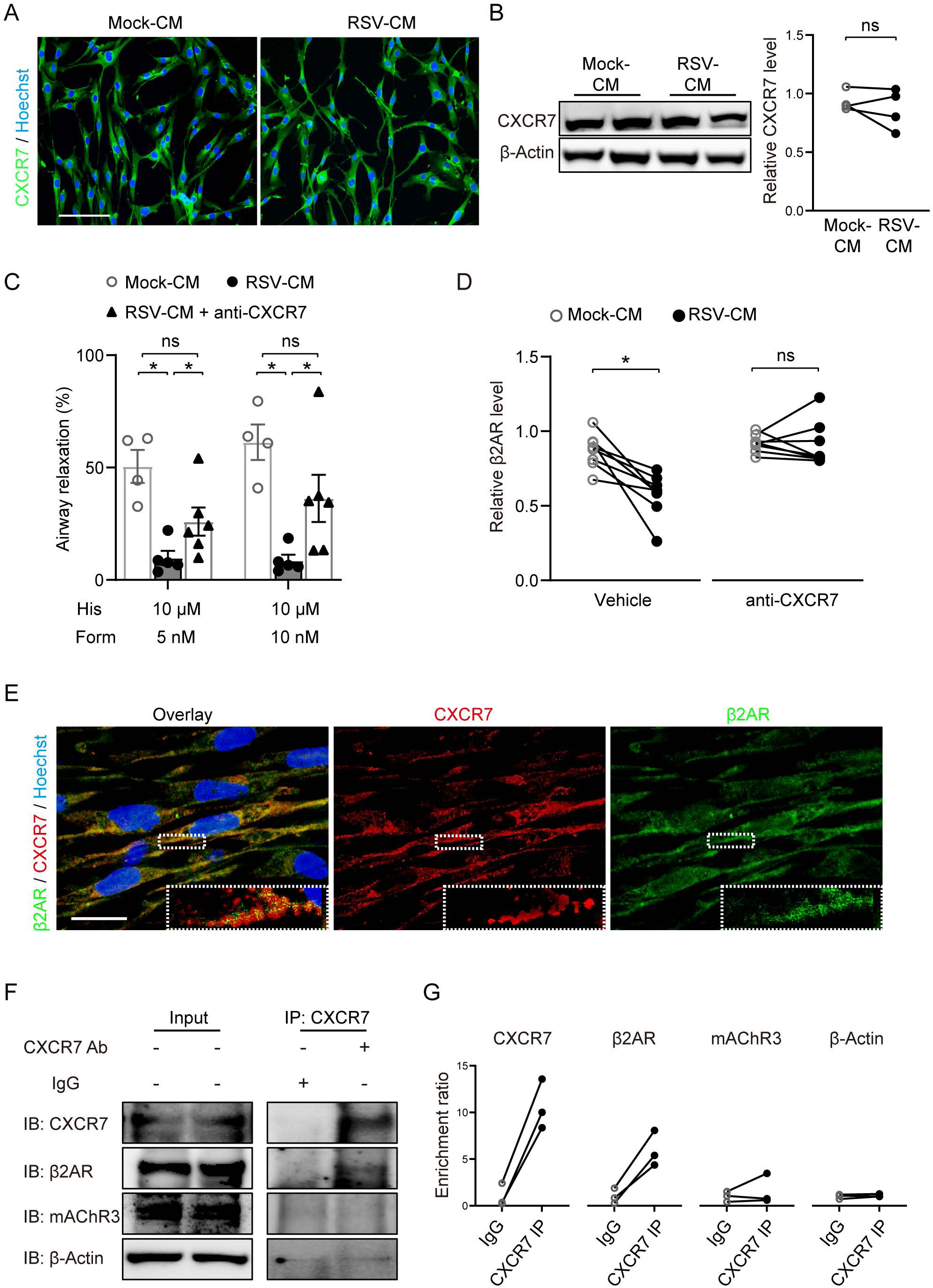
CXCR7 mediates the regulatory role of CXCL11 by interacting with the approximal β2AR in infant ASMCs. (**A**) Representative fluorescence images of CXCR7 staining in infant ASMCs following 48-hr treatment with Mock- or RSV-CM collected from neonatal ALI cultures. Scale bar, 100 μm. (**B**) Representative Western blot for CXCR7 in infant ASMCs after 48-hr treatment with Mock- or RSV-CM of neonatal ALI cultures. β-actin was the loading control in densitometry. N=4 donors. (**C**) Summary plots of 10μM His-precontracted infant airway in response to Form at indicated concentrations in infant PCLSs that were incubated with Mock-CM, RSV-CM, and RSV-CM+CXCR7 antagonist (10 µM) for 48 hours. N=4-6 slices from 4 donors. (**D**) Representative Western blot for β2AR in primary infant ASMCs following 48-hr treatment with Mock- or RSV-CM in the presence of CXCR7 antagonist (10 µM) or vehicle control. β-actin was the loading control in densitometry. N=4 donors with 2 independent repeats. (**E**) Representative confocal fluorescence images showing colocalization of CXCR7 and β2AR on the cell membrane of infant ASMCs. Scale bar,10 μm. (**F**) Representative co-immunoprecipitation (IP) assay for β2AR and mAChR3 by the CXCR7 antibody. β-actin was the loading control for input cell lysates. (**G**) Densitometry to measure the enrichment of CXCR7, β2AR, and mAChR3 by the CXCR7 antibody normalized to IgG in the co-IP assay in (F). Statistical significance was calculated by the Kruskal-Wallis test with a post-hoc test with BYK procedure in (C) and the Wilcoxon test in (B) and (D) *p<0.05. ns, not significant.

CXCR7 has no intracellular domain to activate G protein-coupled signaling and thus is considered a decoy GpCR (33). We thus hypothesized that CXCR7 may cause β2AR desensitization through mechanisms at the cell surface level. Co-staining for CXCR7 and β2AR in ASMCs followed by confocal microscopy showed that these two receptors were detected on the cell surface in close proximity (Fig. 4E). To test whether these two receptors resided in a cell surface complex, we performed pull-down assays using the CXCR7 antibody and detected enrichment of β2AR in the pull-down (Fig. 4F, 4G). In contrast, mAChR3 was not enriched by the CXCR7 antibody (Fig. 4F, 4G). In addition, β-actin, which was loading control for input cell lysates, was barely detectable in the pulldown, validating the efficacy of the CXC7 antibody in enriching cell surface proteins (4F, 4G). These findings suggest that CXCR7 forms a complex with β2AR on the cell surface, which may act to promote β2AR internalization upon binding to CXCL11.

### RSV infection of BALB/c pups impairs β2AR-mediated airway relaxation via the CXCL11-CXCR7 axis

We tested the role of the CXCL11-CXCR7 pathway in β2AR desensitization *in vivo* using RSV-infected BALB/c pups (29, 34). BALB/c pups were infected intranasally with RSV (A2-line 19F, 1×10^6^ PFU) at postnatal day 7 (P7) and P8 and analyzed at P10 for infection and airway physiology (Fig. 5A). As we showed previously (29), RSV-induced epithelial infection and elevated cytokine production in the lung tissues, including *Cxcl11*, in this preclinical BALB/c neonatal model (Fig. 5B, S3). Because mice at P10 are too small to measure airway reactivity by Flexivent, we prepared PCLSs from RSV-infected BALB/c pup lungs for contraction and relaxation assay. We found no difference in airway contraction in response to methacholine up to 0.5 µM; however, PCLSs from RSV-infected BALB/c pups showed blunted airway relaxation in response to formoterol (Fig. 5C). In addition, Cxcr7 neutralization during RSV infection led to a recovery of the relaxation responses to formoterol (Fig. 5D and E). Therefore, the BALB/c neonatal RSV infection model confirms the critical role of Cxcl11-Cxcr7 signaling in β2AR desensitization and intractable airway contraction.

**Fig 5.**
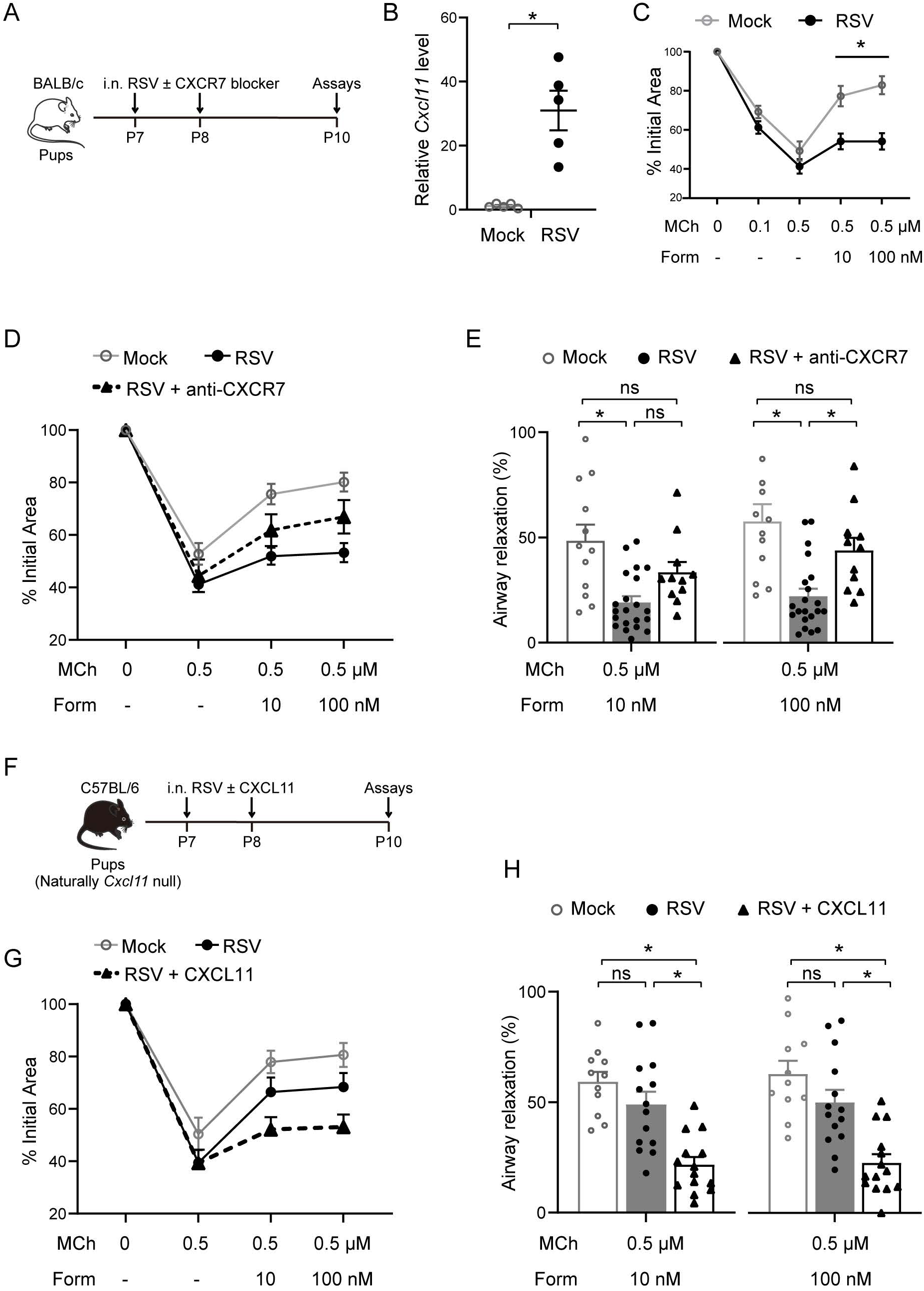
Activation of the CXCL11-CXCR7 pathway downregulates airway relaxation to β2AR agonists in RSV-infected mouse pups. (**A**) Schematic of RSV A2-line19F infection of BALB/c mouse pups at postnatal day 7 (P7) and P8 (1×10^6^ PFU/10 µl) with and without the CXCR7 antagonist (10µM, 10 µl). At day 3 post-infection, mice were analyzed for RSV infection and airway contraction using prepared PCLSs in (B-E). (**B**) *Cxcl11* gene expression in the lungs from Mock- and RSV-infected BALb/c pups by qRT-PCR. Data were normalized to 18s rRNA. Each mark represents one mouse. (**C**) Summary plots of airway contraction in response to methacholine (MCh) and relaxation in response to Form at indicated concentrations. N=8 PCLSs in 3 Mock-infected pups. N=18 PCLSs from 5 RSV-infected pups. (**D**) Summary plots of airway contraction by 0.5 µM MCh and relaxation in response to Form at indicated concentrations. N=12 PCLSs from 4 Mock-infected pups. N=21 PCLSs from 5 RSV-infected pups. N=11 PCLSs from 4 RSV+CXCR7 antagonist pups. (**E**) Airway relaxation in response to 10 and 100 nM Form in (D) was replotted for comparison between the three groups. (**F**) Schematic of RSV A2-line19F infection (1×10^6^ PFU/10 µL) of C57BL/6 mouse pups at P7 and P8 with and without CXCL11 (200 ng/mL, 10 µL). At day 3 post-infection, pups were analyzed for airway contraction using prepared PCLSs (**G**, **H**). (**G**) Summary plots of 0.5 µM MCh-induced contraction followed by 10 and 100 nM Form treatment to induce relaxation. N=11 PCLSs from 3 Mock-infected pups. N=14 PCLSs from 4 RSV-infected pups. N=13 PCLSs from 4 RSV+CXCL11 pups. (**H**) Replotted 10 and 100 nM Form-induced airway relaxation in the three groups. Statistical significance was performed with a two-way ANOVA test with a post-hoc Sidak test in (C), a Mann-Whitney test in (B), and a Kruskal-Wallis test with a post-hoc test with BYK procedure in (E) and (H). *p<0.05, and ns, not significant.

Leveraging the genetic background difference between BALB/c and C57BL/6 strains (the latter is naturally deficient in Cxcl11) (35, 36), we generated a C57BL/6 neonatal model of RSV infection (Fig. 5F) and tested the effect of administered Cxcl11 on airway relaxation in response to β2AR. Notably, previous research has demonstrated that restoration of the Cxcl11 expression in C57BL/6 mice had no effect on virus-induced immune responses (36), probably due to the functional redundancy of CXCL11 with other cytokines, such as CXCL9 and CXCL10 (35, 37, 38). At P10, we found that RSV-infected (A2-line 19F, 1×10^6^ PFU) C57BL/6 pups showed a similar airway relaxation response to formoterol as Mock control (Fig. 5G, 5H). However, when CXCL11 (200ng/ml, 10µl) was delivered together with RSV (Fig. 5F), formoterol-induced airway relaxation was profoundly impaired (Fig. 5G, 5H). Taken together, our findings from neonatal mouse models of RSV infection using *Cxcl11*- competent BALB/c strain and *CXCL11*-null C57BL/6 strain indicate that activation of CXCL11-CXCR7 signaling causes impaired β2AR-mediated ASMC relaxation *in vivo*.

### Nasopharyngeal aspirates from infants with severe RSV bronchiolitis activate the CXCL11-CXCR7 pathway in infant PCLSs rendering the failure of β2AR agonists

To test whether *in vivo* RSV inflammation activates the CXCL11-CXCR7 pathway to cause β2AR desensitization in ASMCs, we collected NPA samples from pediatric patients (2-24 weeks of age) admitted to the intensive care unit (ICU) for severe RSV bronchiolitis (RSV-NPA) and removed RSV by filtering through with a 100 kDa membrane (Fig. 6A). As control, nonRSV-NPA samples from ICU patients (11-36 weeks of age) with rhinovirus or metapneumovirus bronchiolitis was similarly collected and prepared. Of note, nasal epithelial cells can serve as a surrogate to lower respiratory epithelial cells for studies of respiratory infection in young children (39, 40). ELISA of virus-free NPA showed a higher level of CXCL11 in RSV-NPA than in nonRSV-NPA (Fig. 6B). To match the DMEM-F12 medium used for infant PCLS culture, we concentrated virus-free NPA 10 times by centrifugal ultrafiltration using a 3 KDa filter followed by reconstitution to 1 X using the DMEM-F12 medium. The reconstituted virus-free NPA was then used to treat infant PCLSs. After 48 hours, RSV-NPA treatment had no effect on airway contraction in response to histamine but significantly reduced airway relaxation responses to formoterol (Fig. 6C and D). The relaxation response to formoterol was partially restored in the presence of the CXCR7 neutralizing antibody (Fig. 6E and F). In contrast, other cytokines known to be induced by infant RSV infection, such as TNFα (100ng/ml) and IL-1β (10ng/ml), had no significant effect on airway relaxation in infant PCLSs after treatment for 72 hours (Fig. 6G). Therefore, the CXCL11-CXCR7 pathway contributes to the blunted airway relaxation response to β2AR agonists in infant RSV bronchiolitis.

**Fig 6.**
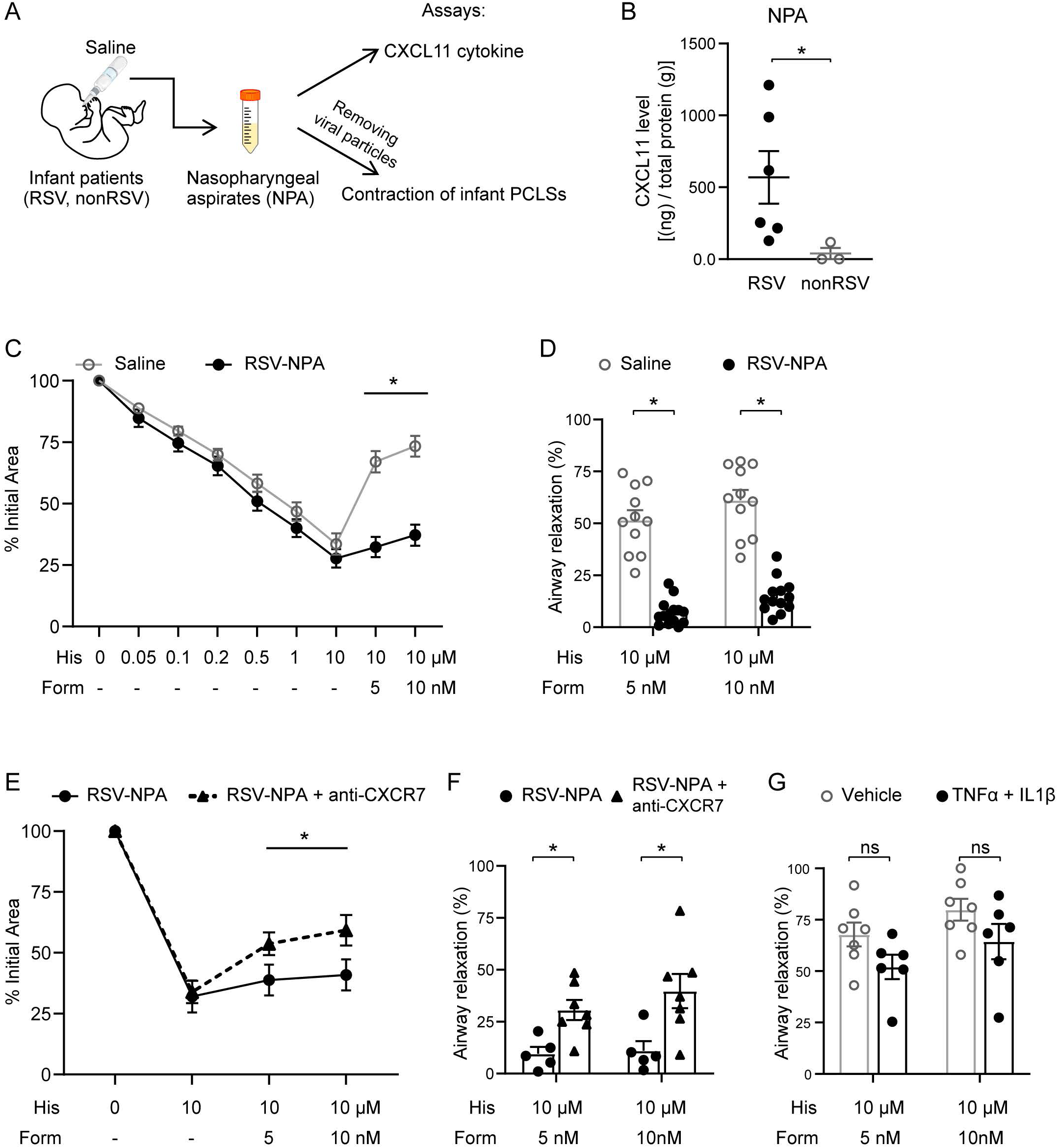
Nasopharyngeal aspirates from infants with severe RSV bronchiolitis suppress β2AR-mediated airway relaxation partially via the CXCL11-CXCR7 pathway. (**A**) Schematic of nasopharyngeal aspirate (NPA) collection from infants with bronchiolitis caused by RSV (RSV-NPA) or by rhinovirus or metapneumovirus (nonRSV-NPA). (**B**) ELISA of CXCL11 levels in RSV-NPA and nonRSV-NPA samples. Each mark represents one patient. (**C**) Summary plots of airway contraction in infant PCLSs in response to His and relaxation in response to Form at indicated concentrations. Infant PCLSs were pretreated with saline or RSV-NPA reconstituted medium for 48 hours. RSV-NPA was pooled from 6 different infant patients. (**D**) Form-induced airway relaxation in saline (N=11 PCLSs from 4 donors) and RSV-NPA (N=14 PCLSs from 4 donors) groups in (C) was replotted for comparison. (**E**) Summary plots of airway contraction in infant PCLSs in response to His and relaxation in response to Form at indicated concentrations. Infant PCLSs were pretreated with RSV-NPA medium or RSV-NPA medium+10µM CXCR7 antagonist for 48 hours. (**F**) Form-induced airway relaxation in RSV-NPA (N=4 PCLSs from 4 donors) or RSV-NPA+CXCR7 antagonist groups (N=7 PCLSs from 4 donors) in (E) was replotted for comparison. (**G**) Summary plots of Form-induced relaxation of His (10 µM)-precontracted airways in infant PCLSs. The infant PCLSs were treated with vehicle (N=7 PCLSs from 3 donors) or TNFα (100 ng/ml) + IL1β (10 ng/ml) (N=6 PCLSs from 3 donors). Statistical significance was performed with a Mann-Whitney test in (B, D, F and G), a two-way ANOVA test with a post-hoc Sidak test in (C) and (E). *p<0.05, and ns, not significant.

## Discussion

This study leveraged the infant PCLS model, an epithelial ALI culture model, and neonatal mouse models of RSV infection to assess the communication between inflammation in infant RSV bronchiolitis and β2AR desensitization in ASMCs. We identified RSV-induced CXCL11 that signals to CXCR7 on ASMCs to enhance β2AR internalization and degradation. The contribution of the CXCL11-CXCR7 pathway to intractable resistance to β2AR agonists was subsequently validated using nasopharyngeal aspirates from infants with severe RSV bronchiolitis as a surrogate to inflammation *in vivo*. All our models of RSV infection allow drug accessibility to ASMCs, as there are no mucus plugs in small airways. In addition, we found no evidence of direct RSV infection of human ASMCs using the human infant PCLS model, which reproduces RSV-induced airway pathophysiology found in patients (29, 41, 42). Therefore, while poor drug accessibility to the affected airways remains a possible cause of therapeutic failure of β2AR agonists in infant RSV bronchiolitis, our findings indicate that β2AR desensitization in ASMCs caused by the inflammatory signal CXCL11 may also contribute to this devastating clinical symptom.

The β2AR-mediated cAMP pathway is the most critical relaxant mechanism operating in ASMCs and has been the focus of bronchodilation therapy. The activity of this pathway is tightly regulated at multiple levels. Our study indicates that RSV inflammation dysregulates cAMP-mediated relaxation at the cell surface β2AR level without affecting the downstream components. Therefore, compounds that activate the intracellular cAMP-PKA pathway could be developed as novel therapeutics to alleviate airway contraction. In addition to CXCL11 in this study, previous work identified other inflammatory factors, such as TNFα and IL-1β, that render β2AR desensitization in human primary ASMC culture, mouse or rabbit tracheal segments (21, 43–45). However, we did not observe a substantial reduction of β2AR-mediated infant airway relaxation following TNFα and IL-1β treatment in our infant PCLS model. We suspect the discrepancy could relate to species-specific regulation of ASMC contraction and relaxation and differences in the age group between this work and previous studies. Although TNFα and IL-1β may not be significant in the relaxant regulation of infant RSV bronchiolitis, our findings of partial recovery of airway relaxation responses to β2AR agonists after blockade of CXCL11 and CXCR7 suggest other inflammatory signals are at play. Future investigations are warranted to identify additional cytokines that may interfere with the β2AR pathway in infant ASMCs in RSV bronchiolitis.

CXCL11 is a small cytokine strongly induced by INFγ and INFβ and generally acts on its receptor CXCR3 on immune cells to recruit them to cancer or inflamed tissues(46, 47). In this study, we uncovered a novel role of CXCL11 in β2-AR desensitization. During in vivo RSV infection, we anticipate CXCL11 to be released from immune cells and epithelial cells (38, 48). CXCL11 executes its role via a decoyed receptor CXCR7 on the cell surface of ASMCs. Due to the lack of G protein binding and signaling, CXCR7 has been traditionally viewed as a scavenger receptor responsible for turnovers of its cognate ligands, CXCL11 and CXCL12 (49). However, a growing body of evidence suggests CXCR7 can also act by interacting with different GpCRs at the cell membrane. For instance, CXCR7 can colocalize with EGFR by using β-arrestin as a scaffold protein (50) or form heteromeric complexes with the α1-adrenergic receptor on vascular smooth muscle(51). Through such interactions, CXCR7 can interfere with the function of heterogeneous receptors. In addition, β2AR is also capable of interacting with cytokine receptors. For example, β2AR on myocardial cells was shown to interact with a chemokine receptor CXCR4 by hetero-dimerization, which led to the downregulation of β2AR-mediated myocardial constriction by CXCR4 ligands (52). Lately, the CXCR4-β2AR hetero-dimers have also been reported in human cancer cells, and the functional significance of these oligomeric complexes is under investigation (53). Consistent with the unconventional role of CXCR7 as a decoy receptor, CXCR7 and β2AR were colocalized on the cell surface of ASMCs. We speculate that the activated CXCR7 may recruit β-arrestin2 and GRK to enhance β2AR phosphorylation and trigger subsequent internalization and degradation. Future studies are required to fully evaluate how CXCR7 and β2AR interact and whether they form a complex so that activation of CXCR7 by CXCL11 can engage β2AR through mediators, such as β-arrestin2.

One major limitation of this study is a lack of clinical lung samples from infants with severe RSV bronchiolitis. However, lung biopsy is contraindicated in the affected infants. In addition, fetal cases of infant RSV disease carry multiple confounding factors, including co-infection and treatment procedures, that may damage ASMCs and affect their β2AR expression. Furthermore, assessment of changes in cell surface β2AR levels is technically difficult using histological sections prepared from clinical lung samples with RSV disease. To overcome this notable technical limitation, we have employed three RSV infection models and analyzed the level, cell surface location, and function of β2AR in ASMCs in this study. Our findings consistently indicate a role of CXCL11 as a critical inflammatory signal in ASMC β2AR desensitization in infant RSV bronchiolitis.

In summary, this study highlights ASMC dysfunction caused by RSV inflammation in refractory airway obstruction in the affected infants and identifies the CXCL11-CXCR7 pathway as a therapeutic target in restoring the bronchodilation effect of β2AR agonists.

## ACKNOWLEDGEMENTS

This work was supported by NIH grants to X.A. (1R01HL154549), P.H.L. (R21AI156597), and Charles Hood Foundation Grant to Y.B. and funds from the Department of Pediatrics at MGH for Lung Cell Bank to X.A.

## AUTHOR CONTRIBUTIONS

C.Z. prepared the infant bronchial epithelial air-liquid interface (ALI) culture and mouse pup models of RSV infection, performed experiments using these modes, and analyzed the bulk RNA-seq data from primary infant airway smooth muscles (ASMCs) upon experimental exposures. A.T. performed the flow cytometry assay of β2AR expression on the cell surface of primary infant ASMCs. E.B. collected the nasopharyngeal aspirates of infant patients from clinical providers after obtaining consent from the patient’s health care proxy. R.F. guided RSV infection experiments. P.H.L. guided the clinical sample acquirement. Y.B. prepared infant PCLSs and performed the experiments using the infant PCLS model of RSV infection and primary ASMCs upon various experimental treatments. X.A. prepared the primary bronchial ASMCs from infant donor lungs. Y.B. and X.A. conceived the study. Y. B. wrote the manuscript. X.A. edited the manuscript. All the authors read and commented on the manuscript.

