## Supplementary information and figures for "Infant RSV infection desensitizes β2-adrenergic receptor via CXCL11-CXCR7 signaling in airway smooth muscle"

#### SUPPLEMENTARY MATERIAL AND METHODS

**RSV Preparation:** The RSV source and preparation in the lab have been described in detail previously. Briefly, RSV strain A2 (VR-1540) was purchased from ATCC. RSV A2-line19F strain was a gift from Dr. Nicholas W Lukacs (University of Michigan). Hep-2 cell line (ATCC CCL-23) was applied to propagate RSV using an established protocol, i.e., Hep-2 cells were infected at 80% confluence with 0.01 multiplicity of infection (MOI) of RSV at 37°C for 1 hour and incubated for 5 days before being scraped into the medium. The supernatant of the medium was collected, aliquoted, and stored at -80°C. RSV virus titers were measured via plaque assay as previously described (1).

**Human precision-cut lung slice (hPCLS) model of RSV infection and airway contraction-relaxation assay**: Human precision-cut lung slices (hPCLSs) from infant donors were prepared and cryopreserved as previously described (2). Frozen hPCLSs were thawed and exposed to RSV (A2 strain, $1x10^6$ PFU) in DMEM/F-12 medium supplemented with anti-anti (Thermo Fisher Scientific, 15240-062) for 72 hours. The RSV infection of the airway epithelial layer was confirmed by detecting intracellular RSV fusion protein in ciliated cells using immunofluorescent staining. For the airway contraction-relaxation assay, hPCLSs were stimulated with histamine followed by formoterol at increasing concentrations. Then, PCLSs were washed with PBS and fixed for 3 hours in 4% PFA, followed by staining and confocal imaging.

**Human epithelial airway-liquid interface culture model of RSV infection:** An age-specific airway epithelial air-liquid interface (ALI) culture was established using basal stem cells derived from tracheal aspirate (TA). The TA samples were collected from neonatal or adult patients intubated for cardiogenic or neurogenic respiratory failure (Massachusetts General Hospital IRB protocol No. 2019P0032960). After 21 days in the ALI culture, airway epithelial cells were fully differentiated and then exposed to RSV ($4X10^5$ PFU in 100-200 μL) or the mock medium on the apical surface following established protocol (3). The basal medium of Mock-(Mock conditioned medium, Mock-CM) or RSV-treated (RSV conditioned medium, RSV-CM) neonatal or adult ALI cultures was collected 48 hours post-infection to simulate the in vivo RSV-induced epithelial inflammation.

**Isolation of primary hASMCs and coculture with RSV-CM from neonatal ALI culture:** Briefly, airway smooth muscle strips were dissected from the main and lobular bronchi of infant donors, followed by the scrape of epithelia using cotton swabs. The muscle strips were then cut into 1-2 mm pieces and placed in the culture plate with the luminal side facing down. The first passage of primary hASMCs (P1) grew out of small pieces and proliferated in the culture plate over 7-14 days to confluence. Subsequently, the primary ASM cells were collected, cryopreserved, or sub-cultured. The first 4 passages were used for experiments. Primary ASMCs were grown in the DMEM/F12 medium supplemented with 10% FBS and anti-anti to 80% confluence and then exposed to the Mock- or RSV-CM from neonatal ALI culture supplemented with 2% FBS (serum reduced medium) for 48 hours. We do not use ITS (insulin-transferrin-selenium) to fully replace FBS due to insulin's impact on β2AR(4).

**Western blot and Immunoprecipitation:** Primary hASMCs were lysed using RIPA buffer supplemented with a complete protease inhibitor cocktail (Sigma Aldrich 11836170001), and phosphatase inhibitor cocktail (EMD Millipore 524625) and then subjected to the western blot assay following standard protocol. Primary antibodies include rabbit anti-β2-adrenergic receptor (1: 500, Abcam ab 182136), mouse anti-β2-adrenergic receptor (1:500, ThermoFisher Scientific MA5-38441), rabbit anti-phosphorylated-β2-adrenergic adrenergic receptor (1:500, ThermoFisher Scientific PA-38403), rabbit-anti CXCR7 antibody (1:1000, ThermoFisher Scientific PA3-069), mouse anti-β-actin (1: 2000; Sigma Aldrich A5441). HRP-conjugated secondary antibodies include goat anti-mouse (1:2000; BD Bioscience 554002) and goat anti-rabbit (1: 1000). For co-immunoprecipitation (Co-IP), 200 μl cell lysate of $10^6$ ASMCs was incubated with 5 μl CXCR7 antibody overnight at 4 ℃ and then adding 20 μl protein G Agarose Beads (Cell signaling 37478) for 2 hours. Immunoprecipitants were recovered by centrifugation, washed with ice-cold lysis buffer, and mixed with protein-loading buffer. Denatured proteins were analyzed by western blot assay. A conformation-specific secondary antibody (Cell signaling 5127) was used to reduce background signaling in Co-IP assay. The antigen-antibody complex was detected by the Supersignal West Dura Extended Duration Substrate (ThermoFisher Scientific 34075). Densitometry measurements were performed with ImageJ.

**Antibody staining:** Primary hASMCs were fixed with 4% paraformaldehyde-PBS for 15 minutes and hPCLSs for 3 hours before immunofluorescent (IF) staining. Antibodies include mouse anti-RSV Fusion protein (1:100,

Bio-Rad MCA490) to identify RSV-infected cells, Cy3-conjugated mouse anti-α-smooth muscle actin (1:100, Sigma-Aldrich C6198) to identify airway smooth muscle cells in hPCLSs, rabbit anti-β2-adrenergic adrenergic receptor, and rabbit anti-muscarinic 3 receptor (1:100, Abcam ab126168). The fluorophore-conjugated secondary antibodies include goat anti-rabbit Alexa Fluor 594 (1:200, ThermoFisher Scientific A-11037) and goat anti-mouse Alexa Fluor 488 (1:200, ThermoFisher Scientific A-11029). Nuclei were stained with Hoechst dye (1:500, ThermoFisher Scientific H3570).

**Transcriptome analysis of primary HASMCs by bulk RNA-seq:** RNA (2 µg per sample) of primary hASMCs ($1x10^6$) was extracted with the Qiagen RNA RNeasy Kit (74104, Qiagen) and sent for Bulk RNA-seq service by Genewiz (South Plainfield, NJ, USA). The Illumina TruSeq V2 Kit (20020594, Illumina) was applied to generate the libraries using Illumina NextSeq 500 as paired-end 42-nt reads. The reads were subsequently mapped to the human hg38 reference genome using STAR algorithm version 2.6.0a. Only genes with an average count above 2 were analyzed. The differential expression and principal component analyses were performed with the DESeq2 R package (1.30.0). Genes with p-values less than 0.05 were identified as differentially expressed genes (DEGs). GSEA was performed with the whole gene list by comparing the RSV-CM group versus the Mock-CM group.

**Gene Expression analysis:** Total RNA was extracted from human primary ASMCs or epithelial ALI culture using an RNeasy kit (QIAGEN, 74106) and was then used to generate cDNA using Superscript III Reverse Transcriptase (ThermoFisher Scientific 18080-044). Real-time PCR was performed with CFX96 real-time system (Bio-Red) using SYBR Green Mix (ThermoFisher Scientific 4367659). The relative level of gene expression was measured by normalizing to *18S* rRNA using ΔΔCt (cycle threshold difference). All primers were listed in Table S2.

**Cytokine quantification:** The Mock- and RSV-CM from neonatal and adult epithelial ALI cultures were collected. The RSV-induced epithelial cytokines and chemokines were assessed using a commercial kit (Proteome Profiler Human XL Cytokine Array Kit, R&D Systems ARY022B). The amount of CXCL11 in the medium was measured using an ELISA kit (Human CXCL11/I-TAC ELISA Kit, R&D Systems DCX110) following instructions.

**Reconstitution of culture medium using Nasopharyngeal Aspiration from infants with RSV bronchiolitis:** Nasopharyngeal aspiration (NPA) was performed by nursing staff in the pediatric ICU following the protocol; ~3 ml sterile saline lavage solution was acquired each time. The supernatant was collected, aliquoted, and stored at -80 °C. During experiments, we thawed one aliquot (1ml) from each donor and mixed them before processing. Then, we removed viral particles from NPA samples with centrifugal ultrafiltration (Amicon 100kDa, Millipore Sigma). The RSV-free flowthrough will be concentrated to 10X using ultrafiltration (Amicon 3kDa) and then reconstituted to 1X solution using DMEM/F-12 culture medium, called NPA medium. The control medium was prepared by adding normal saline to the DEM/F-12 medium at a 1:9 volume ratio. Freshly prepared control or NPA medium was applied to treat infant PCLSs.

**Quantification of β2AR expression on airway smooth muscle cell surface:** We used 2 approaches, flow cytometry and cell-surface protein biotinylation followed by avidin isolation and a western blot assay, to quantify the β2AR expression on the cell surface of hASMCs. After experimental treatment, live hASMCs were collected into Eppendorf tubes ($1.2 \times 10^6$ per vial). For flow cytometry assay, cells were incubated with β2AR antibody (1:100) for 1 hour, washed with PBS, and then incubated with goat anti-mouse Alexa Fluor 488 (1:100) for 30 mins before detection with a flow cytometer. All staining steps were performed at 4°C. The control hASMCs were only stained with fluorophore-conjugated 2$^{nd}$ antibody. After staining, cells were analyzed on Cytek Aurora. Flowjo v.10.10 was used for data analysis. For the biotinylation assay, cells were incubated with 0.8μM Sulfo-NHS-SS-biotin (ThermoFisher Scientific 21331) PBS solution for 30 mins on ice, washed with ice-cold quenching solution (PBS with 100mM glycine), lysed with 1x lysis buffer (120 μL/vial) containing protease inhibitor for 30 mins on ice. The cell lysate was centrifuged (20min, 14000 rpm, 4°C), and the supernatant was collected. A small volume (20 μL) was taken out for total protein quantification. The rest of the volume was incubated with Neutravidin agarose resin (50 μL, ThermoFisher Scientific 29204) overnight at 4°C to isolate the biotin-labeled surface protein. After centrifuge, supernatant removal, and wash, the agarose resin was dissolved in 1x lysis-LDS buffer to a final volume of 80μL and denatured at 65°C. 40 μL denatured agarose was applied to detect the β2AR expression with western blot assay. The β-actin of the total cell lysate supernatant (20 μL) was used as a control, reflecting the total protein in each sample.

**Table S1. Demographic information of infant patients with viral infections.**

| **RSV-NPA** | | |
|---|---|---|
| Patient Study ID | Age | Viruses |
| RSV1 | 2 weeks | Respiratory syncytial virus |
| RSV2 | 8 weeks | Respiratory syncytial virus |
| RSV3 | 16 weeks | Respiratory syncytial virus |
| RSV4 | 24 weeks | Respiratory syncytial virus |
| RSV5 | 9 weeks | Respiratory syncytial virus |
| RSV6 | 4 weeks | Respiratory syncytial virus |
| **NonRSV-NPA** | | |
| Patient Study ID | Age | Viruses |
| NonRSV1 | 11 weeks | Rhinovirus |
| NonRSV2 | 28 weeks | Metapneumovirus |
| NonRSV3 | 36 weeks | Rhinovirus |

**Table S2. List of primers used for RT-qPCR.**

| **Gene** | **Forward Primer** | **Reverse Primer** |
|---|---|---|
| Human *GAPDH* | GGAGCGAGATCCCTCCAAAAT | GGCTGTTGTCATACTTCTCATGG |
| Human *CXCR7* | GAAGAGATGCAGATCCATCGT | GCTCACAGTTGTTGCAAAGTG |
| Human *CXCR3* | CTCGGCGTCATTTAGCACTT | CTCGGCGTCATTTAGCACTT |
| Mouse *18s* | CCATTCGAACGTCTGCCCTAT | GTCACCCGTGGTCACCATG |
| Mouse *Cxcl10* | CCGGAATCTAAGACCATCAAG | GAGGCTCTCTGCTGTCCATC |
| Mouse *Cxcl11* | CTGGGATTCACCTCAAGAACATC | CAGGGTCAAGGCAAGCCTC |
| Mouse *Il6* | GGCCTTCCCTACTTCACAAG | ATTTCCACGATTTCCCAGAG |
| Mouse *Tnf* | CCCTCACACTCAGATCATCTTCT | GCTACGACGTGGGCTACAG |
| Mouse *Ifna* | CTTCCACAGGATCACTGTGTACCT | TTCTGCTCTGACCACCTCCC |
| Mouse *Ifnb1* | AGATCAACCTCACCTACAGG | TCAGAAACACTGTCTGCTGG |
| Mouse *Ccl2* | TTAAAAACCTGGATCGGAACCAA | GCATTAGCTTCAGATTTACGGGT |

#### SUPPLEMENTARY FIGURE LEGENDS

**Fig S1. RSV-induced epithelial inflammation modified the mRNA profile of infant ASMCs.**

(**A**) Schematic of bulk RNA-seq of infant ASMCs (n = 3 donors) with treatment of Mock-CM or RSV-CM.

(**B**) Volcano plot showing differentially expressed genes.

(**C**) Heatmap showing changes of selected genes related to the relaxant regulation.

(**D**) GSEA enrichment plot showing top pathways in RSV-CM-treated ASMCs versus Mock-CM-treated ASMCs.

**Fig S2. RSV induced age-related epithelial inflammation and impact on infant airway relaxation.**

(**A**) Schematic of cytokine array measurement of conditioned medium.

(**B**) Representative images showing cytokine profiles of the Mock-CM or RSV-CM of neonatal or adult ALI cultures.

(**C**) Schematic of infecting adult epithelium in ALI culture without or with RSV (MOI 2, $4X10^5$) for 48 hours, collecting mock conditioned medium (Mock-CM) or RSV-CM from basal wells, and then applying them to infant PCLSs for 48 hours.

(**D**) Summary plot of Histamine-induced contraction followed by Formoterol-induced relaxation at indicated concentrations in infant airways exposed to Mock-CM or RSV-CM collected from adult ALI culture. N = 8-10 airways from 4 infant donors. ns, not significant by a two-way ANOVA test with a post-hoc Sidak test.

**Fig S3. RSV induced significant lung inflammations in BALb/c pups.**

(**A**) A representative image of double fluorescence staining for a ciliated cell marker (AceTUB) and RSV F protein in the tracheal epithelium of RSV-infected BALB/c pups. Scale bar, 100 μm.

(**B**) The mRNA expression of selected cytokines in the lung lobes of BALb/c pups 72 hours after RSV infection. *$p<0.05$ by the Mann-Whitney U test.

### Figure S1

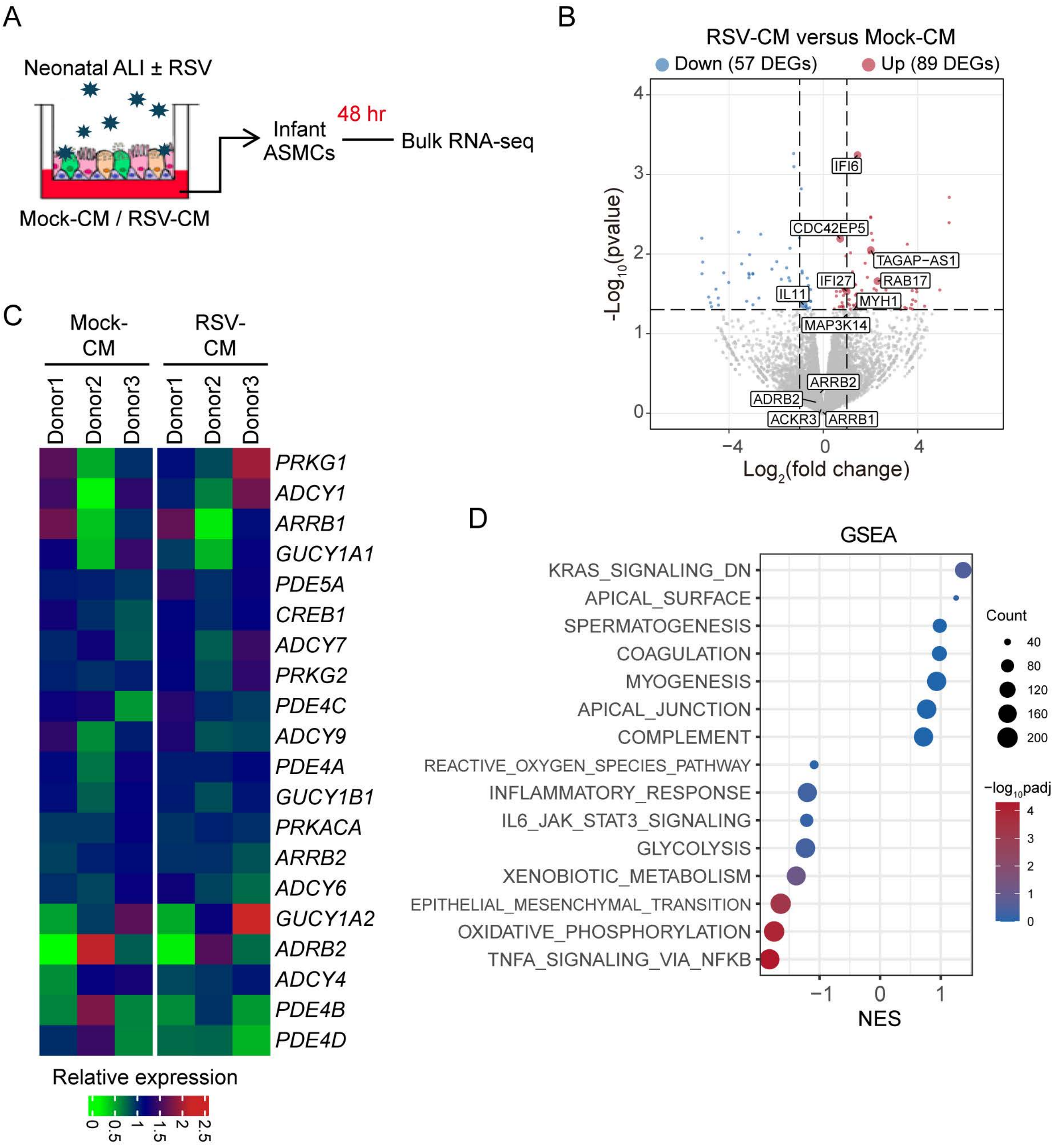
A
Neonatal ALI ± RSV
Mock-CM / RSV-CM
Infant ASMCs
48 hr
Bulk RNA-seq
B
RSV-CM versus Mock-CM
Down (57 DEGs)
Up (89 DEGs)
-Log10(pvalue)
Log2(fold change)
IFI6
CDC42EP5
TAGAP-AS1
RAB17
IFI27
IL11
MYH1
MAP3K14
ARRB2
ADRB2
ACKR3
ARRB1
C
Mock-CM
RSV-CM
Donor1
Donor2
Donor3
PRKG1
ADCY1
ARRB1
GUCY1A1
PDE5A
CREB1
ADCY7
PRKG2
PDE4C
ADCY9
PDE4A
GUCY1B1
PRKACA
ARRB2
ADCY6
GUCY1A2
ADRB2
ADCY4
PDE4B
PDE4D
Relative expression
D
GSEA
KRAS_SIGNALING_DN
APICAL_SURFACE
SPERMATOGENESIS
COAGULATION
MYOGENESIS
APICAL_JUNCTION
COMPLEMENT
REACTIVE_OXYGEN_SPECIES_PATHWAY
INFLAMMATORY_RESPONSE
IL6_JAK_STAT3_SIGNALING
GLYCOLYSIS
XENOBIOTIC_METABOLISM
EPITHELIAL_MESENCHYMAL_TRANSITION
OXIDATIVE_PHOSPHORYLATION
TNFA_SIGNALING_VIA_NFKB
NES
Count
−log10padj

### Figure S2

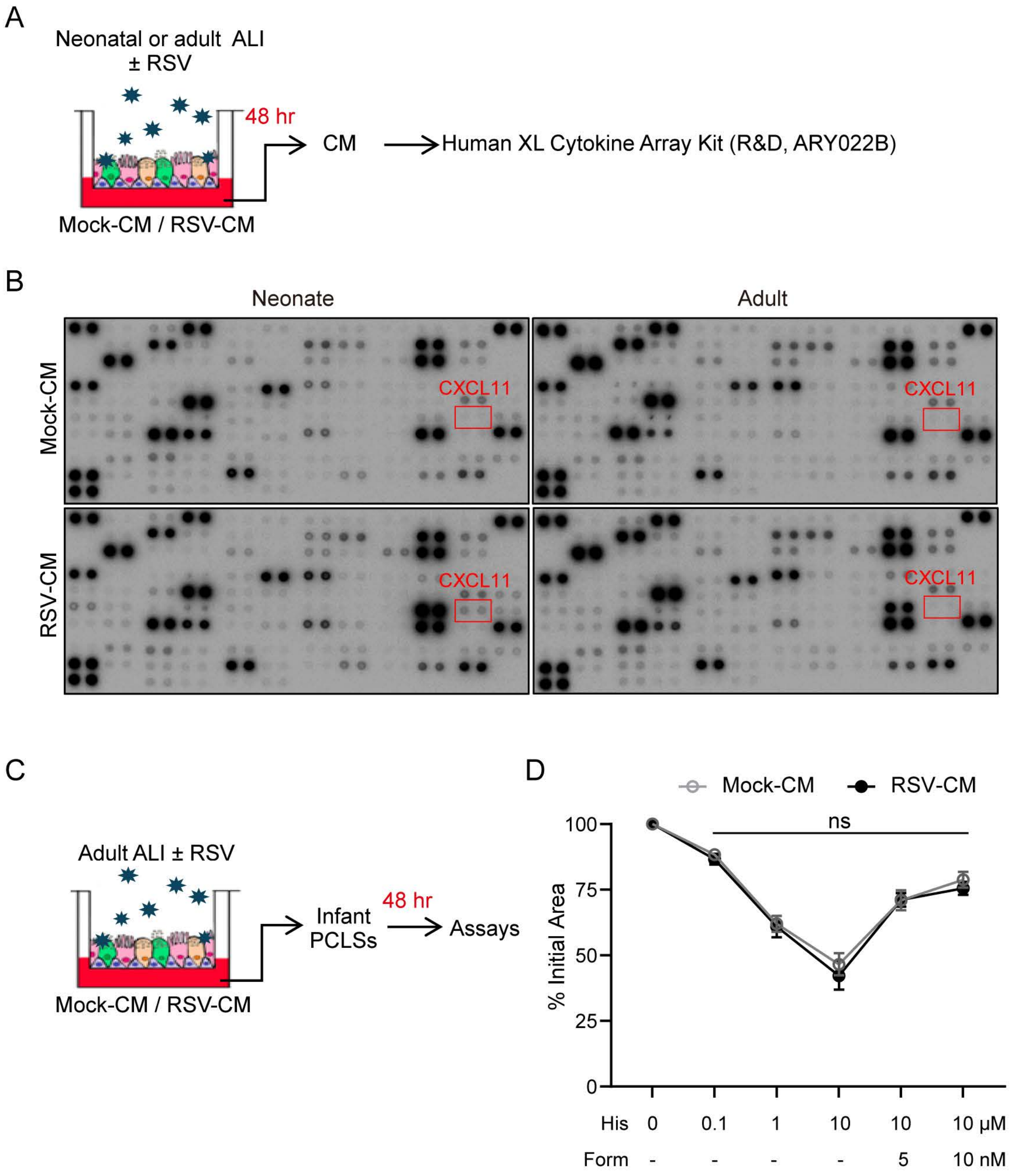

A
Neonatal or adult ALI
± RSV
48 hr
CM
Human XL Cytokine Array Kit (R&D, ARY022B)
Mock-CM / RSV-CM
B
Neonate
Adult
Mock-CM
RSV-CM
CXCL11
C
Adult ALI ± RSV
48 hr
Infant
PCLSs
Assays
Mock-CM / RSV-CM
D
Mock-CM
RSV-CM
ns
% Initial Area
100
75
50
25
0
His 0 0.1 1 10 10 10 μM
Form - - - - 5 10 nM

### Figure S3

A

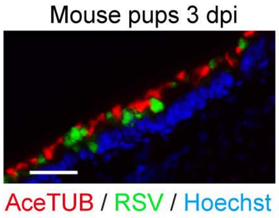
Mouse pups 3 dpi
AceTUB / RSV / Hoechst

B

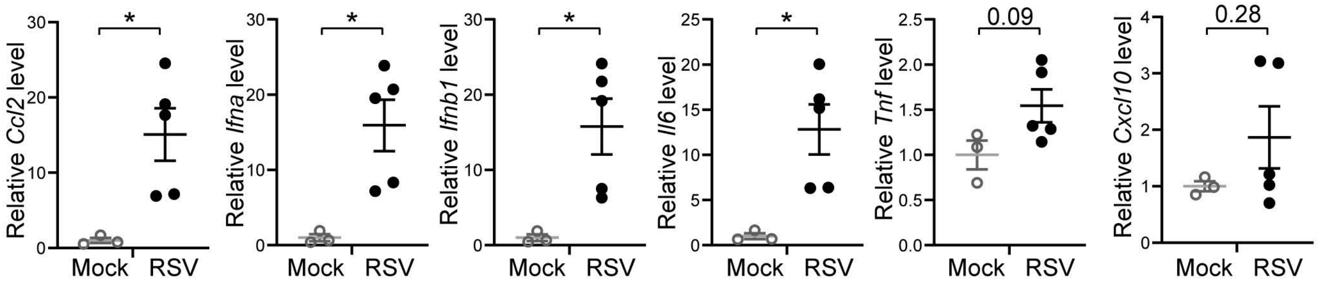
Relative Ccl2 level
Relative Ifna level
Relative Ifnb1 level
Relative Il6 level
Relative Tnf level
Relative Cxcl10 level
*
0.09
0.28
Mock
RSV
